# The Molecular Programme of the Biphasic Isopod Moult: A Transcriptomic Chimera

**DOI:** 10.64898/2026.08.02.742273

**Authors:** Idan Sheizaf, Robert M. Waterhouse, Marc Robinson-Rechavi, Ariel D. Chipman

## Abstract

Isopods are an order of crustaceans characterised by a biphasic moulting pattern, in which the posterior cuticle is shed before the anterior cuticle, with an intramoult period of up to a few days between the two. In order to understand how this unusual moulting pattern is regulated, we carried out a transcriptomic analysis covering three distantly related terrestrial isopod species. We analysed the transcriptomic profile of four body regions: the front legs, the hind legs, the thorax and the head, at different phases of the moulting cycle in the three species. We describe a conserved cyclic pattern in the transcriptomic profiles corresponding to the phases of the moulting cycle. The genes driving this conserved pattern provide a catalogue of the central players of the moulting process and are prime candidates for future experimental work. Furthermore, we show that during the intramoult phase, the posterior limbs display a transcriptomic profile more similar to the postmoult phase, indicating that at this phase, the animal is functionally a transcriptomic chimera, with the anterior and posterior halves experiencing radically different molecular environments, with disjunct regulatory programmes active in each half.

## Introduction

The crustacean order Isopoda exhibits remarkable ecological plasticity, with species occupying diverse habitats, ranging from the abyssal benthos to semi-arid terrestrial environments^1–3^. Within Isopoda, the suborder Oniscidea constitutes the most species-rich group of terrestrial crustaceans, representing a monophyletic lineage that arose from a single successful terrestrialization event^3,4^. This transition from aquatic to land-based life necessitated substantial physiological and morphological adaptations to manage desiccation, gas exchange, and nutrient cycling^5,6^. As primary detritivores, oniscideans play a key role in terrestrial ecosystems by facilitating the decomposition of organic matter^7^.

A defining characteristic of all arthropods is the periodic shedding of the exoskeleton, a process known as ecdysis or moulting, which is required for somatic growth and development^8^. In most arthropods, including crustaceans, the moult cycle can be subdivided into three primary physiological phases^8–13^ The first is the *intermoult* phase, in which the animal is physiologically stable and accumulates mass^10,14^. The preparatory *premoult* phase involves the synthesis of regulatory hormones, calcium resorption, and the biosynthesis of the new cuticular layers^15,16^. At the end of this phase, the animal sheds its old exoskeleton in a relatively short period of time, between minutes and a few hours. Different arthropods have varied mechanisms for exuviation - some, like spiders break through the dorsal part of their exoskeleton^17,18^, others like chilopods break through the anterior while leaving a crumpled exuvia behind^18,19^. Finally, in the *postmoult* phase, the animal’s body expands and hardening of the exoskeleton occurs through mineralization or tanning^20,21^. Some animals, like insects and spiders, moult a specific number of times and reach a final developmental stage where they attain sexual maturity, known as the ‘terminal moult’^17,22^, while others, like most non-insect crustaceans, continue growing and moulting beyond sexual maturity, and possibly throughout their lives^23–26^.

In contrast to the synchronous whole-body moulting observed in other crustaceans, the entire order Isopoda employs a uniquely modified biphasic moulting strategy^6^. Instead of a single exuviation event, isopods shed their exoskeleton in two discrete stages separated by an interval of 24 to 48 hours^27–29^. The process begins with the shedding of the posterior exoskeleton, followed by an intramoult period, and concludes with the shedding of the anterior exoskeleton. Consequently, the isopod moult cycle comprises four distinct stages: intermoult, premoult, intramoult, and postmoult^6,27–29^.

The mechanism underlying this biphasic strategy represents a singular physiological and regulatory challenge, as it requires a precise anatomical decoupling of the organism. We suggest that during the intramoult interval, the posterior half transitions to a post-ecdysial state of hardening, while the anterior half remains in a pre-ecdysial state of cuticle digestion and preparation. This staggering of the moult is a specialized evolutionary adaptation that allows isopods to maintain partial mobility and avoid the complete physical vulnerability associated with simultaneous whole-body exuviation^30^.

At the molecular level, the arthropod moult cycle is governed by a hierarchical endocrine axis and a coordinated series of transcriptomic waves^8^. The cycle is primarily driven by pulses of ecdysteroids (e.g., 20-hydroxyecdysone)^31,32^, which trigger a conserved cascade of gene expression often described by the ‘Ashburner model’^8,33^. In this framework, ecdysteroids first activate a small set of ‘early-response’ transcription factors - such as *E74*, *E75*, and *Broad-Complex* - which in turn orchestrate the activation of ‘late-response’ genes responsible for structural components, including cuticular proteins (CPs) and chitin-modifying enzymes^8,22,31,32,34^. In crustaceans, this process is further modulated by the sesquiterpenoid methyl farnesoate, (MF, the crustacean juvenile hormone analogue), which acts as a regulatory ‘status quo’ switch during development^8,31^. For isopods, we postulate that the biphasic strategy necessitates a distinct spatial and temporal partitioning of these molecular cascades. The organism must coordinate these transcriptomic waves across its anatomical regions, maintaining a late-phase mineralization signature in the posterior half while the anterior half remains under the control of early-phase degradative and preparatory programmes.

To investigate these conserved patterns, we applied a comparative approach across three representative terrestrial isopod species: *Porcellio laevis* (swift woodlouse), *Armadillidium maculatum* (zebra pillbug), and *Armadillo officinalis* (hissing isopod). These species represent three distinct families within the suborder Oniscidea^4^, reflecting diverse ecological niches, behaviours, and morphological adaptations^5,35,36^, with more than 100 My of divergenceֱֱ^37,38^. *P. laevis* (family Porcellionidae) is a widely distributed^39^, fast-running generalist that does not roll into a ball (non-conglobating); it is typically found under rocks and leaf litter, exhibiting relatively high water loss rates and relying heavily on microclimate selection^36^. In contrast, both *A. maculatum* (family Armadillidiidae) and *A. officinalis* (family Armadillidae) are conglobating species that can roll into a sphere to protect their soft ventral surfaces from desiccation and predation^36^. *A. maculatum* is native to Southern France and Monaco in temperate habitats, known for its distinctive black-and-white disruptive striping^40^. *A. officinalis* is a larger species adapted to drier, warmer Mediterranean climates, such as the southern Levant, and is notable for its ability to roll tightly and emit stridulatory sounds^35^. While both *A. officinalis* and *A. maculatum* are conglobating species, they are phylogenetically distinct^4^; the conglobating feature has evolved convergently^41^, and *A. maculatum* is more closely related to *P. laevis* than it is to *A. officinalis*^4^. By examining these three phylogenetically divergent representatives, we can identify robust transcriptomic themes and regulatory programmes that are shared across diverse terrestrial isopods.

In this study, we characterized the comparative transcriptomics of the terrestrial isopod moult cycle by analysing body part-specific expression patterns across *P. laevis*, *A. maculatum*, and *A. officinalis*. We performed RNA sequencing on four distinct body parts (anterior limbs [front legs], posterior limbs [hind legs], heads, and thoraces) collected from animals at four key stages of the moult cycle: intermoult, premoult, intramoult, and postmoult. Using de novo transcriptome assemblies, we mapped transcripts to orthologous groups (OGs) to build a unified comparative expression matrix. We then quantified the global transcriptomic dynamics across phases, analyzed anatomical desynchrony during the biphasic switch (Intramoult) by contrasting anterior and posterior body parts, and identified a core ‘molecular toolkit’ of synchronized biomarkers. By tracing these transcriptomic waves, this work provides a detailed functional genomics framework to understand the evolutionary stability of the biphasic moult programme and its role in supporting the physiological resilience of isopods in terrestrial environments.

## Methods

### Animal Husbandry

Individuals of *A. maculatum* and *P. laevis* (orange strain) were obtained from hobbyists. Individuals of *A. officinalis* were collected in a number of locations in and around Jerusalem. Laboratory colonies of *A. maculatum*, *P. laevis*, and *A. officinalis* were maintained in ventilated plastic containers. Each container was supplied with organic soil and a layer of dried oak leaves to provide both a naturalistic substrate and a primary dietary source. High relative humidity was maintained through regular misting with deionized water. All animals were housed in a climate-controlled environment at 21°C under a 14:10 hour light:dark photoperiod. Moult progression was monitored through daily visual inspection of behavioural changes, cuticle opacity, and the appearance of ventral white patches (calcium carbonate stores) to ensure accurate, stage-specific sampling.

### RNA Extraction and Library Preparation

Total RNA was extracted from four distinct body parts (anterior limbs, posterior limbs, heads, and thoraces) of *A. maculatum*, *P. laevis*, and *A. officinalis* across four key phases of the moult cycle (intermoult, premoult, intramoult, and postmoult). Samples were homogenized and total RNA was extracted using the ZYMO RESEARCH Quick-RNA MiniPrep kit (Cat. No. R1055) following the manufacturer’s protocol. The quality and concentration of each RNA sample were assessed using an Agilent 2100 Bioanalyzer (Eukaryote Total RNA Nano assay, version 2.6); samples showing significant degradation were excluded.

For reference transcriptome construction (*de novo* assembly), libraries were prepared on a Perkin Elmer Sciclone liquid handling robot using 250 ng of RNA input. Library preparation utilized the Illumina Stranded mRNA Kit, incorporating Illumina PolyACapture (ref 20040893), Illumina cDNA Synthesis (ref 20040895), and Illumina RNA Prep, Ligation (ref 20040897), with indexing performed via the IDT for Illumina RNA UD Indexes Set B Ligation (ref 20040553-20040556).

For differential gene expression analysis of phase-specific samples, libraries were prepared on a Perkin Elmer Sciclone liquid handling robot using 400 ng of RNA input. Library preparation utilized the Watchmaker mRNA Library Prep Kit (7BK0001-096), incorporating the Watchmaker mRNA Capture Kit (7K0105-096), the Watchmaker RNA Library Prep Kit (7K0078-096), and the Twist Universal Adapter System - Truseq Compatible 96 sample Plate A (Ref: 101308).

### RNA Sequencing

Sequencing was performed on the Element Biosciences AVITI platform using HD Freestyle flow cells. Libraries prepared for *de novo* transcriptome assembly were sequenced in 2x150 bp paired-end mode using the AVITI 2x150 Sequencing Kit Cloudbreak FS High Output (#860-00013) to obtain deep sequence coverage and facilitate contig resolution. Libraries prepared for differential gene expression were sequenced in single read 150 mode using the AVITI 2x75 Sequencing Kit Cloudbreak FS High Output (#860-00015). Raw base-call files were converted to FastQ format and demultiplexed.

### *De Novo* Transcriptome Assembly and Refinement

Despite initial attempts to sequence and assemble high-quality reference genomes for *A. maculatum*, *P. laevis*, and *A. officinalis*, these efforts were unsuccessful due to high genome size, repetitiveness, and structural complexity common in several groups of crustaceans. Consequently, we focused our efforts on constructing de novo transcriptome assemblies for each species. This approach proved to be highly effective for characterizing relative expression changes and for identifying functional protein-coding transcripts across the distinct stages of the moult cycle.

Raw sequencing reads were first evaluated for quality using FastQC (v0.12.1)^42^. To ensure high-quality data for assembly, adapter sequences and low-quality bases were removed using fastp (v0.23.4)^43^. For single-end reads, default filtering parameters were applied; for paired-end reads, the –detect_adapter_for_pe flag was utilized to ensure synchronized trimming of both directions.

*De novo* transcriptome assembly was performed using Trinity (v2.15.2)^44^ via an Apptainer containerized environment. Assemblies were generated using synchronized paired-end reads with a minimum contig length of 200 bp. To facilitate downstream multi-species comparison, all Trinity FASTA headers were standardized by replacing the default “TRINITY” prefix with species-specific identifiers.

Candidate coding regions (ORFs) were predicted using the TD2 (TransDecoder v2)^45^ suite. To ensure high sensitivity for conserved and taxonomically relevant sequences, an initial set of potential ORFs was generated using TD2.LongOrfs. These candidate regions were subjected to homology searches using MMseqs2 (v15.6f4ea)^46^ against Swiss-Prot^47^, UniRef90^48^, Pfam^49^, and the CrusTome database^50^. High-confidence homology hits - specifically those identified in the crustacean-specific CrusTome database - were used to prioritize and retain ORFs during the TD2.Predict phase, ensuring the preservation of conserved isopod genes.

To reduce transcriptomic redundancy and collapse highly similar isoforms, the predicted coding sequences (CDS) were clustered using CD-HIT (v4.8.1)^51^. A sequence identity threshold of 95% (-c 0.95) was applied to generate a non-redundant representative set of transcripts for each species.

To ensure a high-purity isopod transcriptome, a comprehensive taxonomic filtering pipeline was implemented using MMseqs2 taxonomy. Clustered transcripts were searched against the UniRef90 database to assign taxonomic lineages. Contaminants - specifically sequences originating from Bacteria (taxid 2), Archaea (taxid 2157), Fungi (taxid 4751), Viridiplantae (taxid 33090), and Viruses (taxid 10239) - were identified and removed. Sequences that were either properly classified as Metazoan or remained “Unclassified” (representing potentially novel or highly divergent isopod-specific genes) were retained in the final “Clean” assembly.

Assembly completeness and biological representation were assessed using BUSCO (v5.7.1)^52^ against the crustacea_odb12 lineage. Completeness scores were monitored both before and after taxonomic decontamination to ensure that the filtering process maintained high biological representation while removing non-target sequences.

### Functional Annotation and Orthology Mapping

Functional assignments for the predicted proteomes were primarily derived using EggNOG-mapper (v2.1.12)^53^ against the EggNOG database^54^, providing Gene Ontology (GO) terms, COG categories, and KEGG Orthology (KO) identifiers.

Additionally, signal peptides were predicted using SignalP (v6.0)^55^ to identify putative secreted proteins involved in the moult process.

To ensure high-quality functional representation, KEGG KO identifiers were used to systematically annotate additional GO terms via a custom KO-to-GO enrichment pipeline. This process utilized the KEGG API to map KO numbers to their corresponding biological process (BP) and molecular function (MF) terms, creating a comprehensive annotation master file for each species.

To enable direct comparison of transcriptomic responses across the three species *(A. maculatum*, *P. laevis*, and *A. officinalis*), transcripts were assigned to Orthologous Groups (OGs) using Orthologer ODB-mapper (v3.9.0)^56^.

Representative transcripts from each assembly were mapped against the Crustacea (taxid 6657) OrthoDB reference set (38861 v12.2 Crustacea-level OGs comprising 30 species), assigning each sequence to its most likely ancestral OG (ODB_OG assignment).

This mapping allowed for the integration of disparate species datasets into a single “Universal Isopod Moult Matrix”. Specifically, transcript-level raw counts for each species were first aggregated by their corresponding OrthoDB ODB_OG identifiers, and the resulting species-specific matrices were merged using an inner-join to isolate only the OGs shared across the three species. For genes that did not have an ODB_OG assignment but showed high sequence similarity and shared functional annotations according to EggNOG-mapper, a manually curated “Preferred Name” mapping was used to ensure their inclusion in the final cross-species synthesis.

### Transcript Quantification and Differential Expression

Transcript abundance was quantified using Salmon (v1.10.1)^57^ in mapping-based mode. An index was first constructed from the filtered, non-redundant “Clean” assembly for each species. Scaled reads were mapped to the index using the --validateMappings flag. To account for potential biases in the sequencing data, both sequence-specific and fragment-level GC-bias corrections (--seqBias, --gcBias) were applied. For single-end libraries, a mean fragment length of 250 bp (SD = 25 bp) was specified. Quantification was performed with 100 bootstrap replicates to estimate technical variance.

Salmon quantification files (quant.sf) were imported into R (v4.4.1)^58^ using the tximport package^59^. Differential expression analysis was performed using DESeq2 (v1.44.0)^60^. To identify biologically relevant changes, a multi-factor design was implemented using the formula ∼group, where “group” represented the interaction between body parts and moult phase.

Genes with low counts (total sum < 10) were filtered out prior to analysis. Dispersions were estimated and a negative binomial generalized linear model was fitted to the data. Multiple types of contrasts were performed to capture the full complexity of the moult cycle:

- Temporal Waves: Phase-*vs*-phase comparisons (e.g., premoult *vs*. intermoult) were conducted both across all body parts and within specific body parts.
- Anatomical Asymmetry: Within-phase comparisons between front and back legs were used to identify the “Biphasic Asymmetry” (e.g., Front *vs*. Back legs during intramoult).
- Body part Specificity: Each body part was compared against all others within a given phase to identify body part-specific drivers.

P-values were adjusted for multiple testing using the Benjamini-Hochberg procedure, and genes with an adjusted p-value < 0.05 and an absolute log2 fold change > 1 were considered significantly differentially expressed.

For PCA and heatmap visualizations, raw counts were transformed using the Variance Stabilizing Transformation (VST) provided by DESeq2, which accounts for the mean-variance relationship in high-throughput sequencing data.

### Comparative Analysis and Cross-Species Synthesis

To characterize internal transcriptomic variance within *A. maculatum, P. laevis,* and *A. officinalis*, Principal Component Analysis (PCA) was performed independently on each VST-normalized count matrix. To optimize the resolution of the major variance axes, a range of transcript selection thresholds (top 500, 1000, and 2000 most variable transcripts) was systematically evaluated for each species. To rigorously validate the biological relevance of the resulting clustering, Permutational Multivariate Analysis of Variance (PERMANOVA) was performed using the vegan package (999 permutations) to quantify the effect size (R²) and statistical significance (p-value) of the moult phase on the global transcriptomic state. The threshold providing the most stable representation and highest statistical support for physiological stages (2000 genes) was selected for downstream species-specific visualizations.

To identify conserved transcriptional trajectories across the study species, OGs shared across all three species were used to integrate data from all three species. For OGs represented by multiple transcripts within a single species, raw counts were summed to generate a single value per OG per sample. Comparative analysis indicated that the hind leg sample (*h.legs*) displays a significantly different transcriptomic signal to all other body parts. Therefore, to minimize technical noise and focus on the universal physiological signal, these samples were excluded from the global phase-engine analysis.

To resolve cross-species batch effects (primarily driven by species identity), ComBat-seq (from the sva package^61^) was applied to the integrated raw count matrix, using species as the batch factor and moult phase as the preserved biological signal. VST normalization was subsequently performed on the corrected counts. This approach successfully shifted the primary biological variance (the “Universal Moult Phase”) to the first two Principal Components (PC1 and PC2).

The top 100 OGs driving these conserved axes were extracted based on their absolute feature loadings. To prioritize high-impact candidates, OGs were ranked by an Importance Score, calculated as the product of the absolute loading and the percentage of variance explained by the respective PC. Kruskal-Wallis tests were applied to the Principal Component scores to confirm their significant correlation with the moult phase (p < 0.05).

The conserved transcriptomic response was identified by extracting the overlapping subset of significantly regulated OGs across all three species for three major transitions: premoult, intramoult, and postmoult versus an intermoult reference. Statistical significance for each species was defined as an adjusted p-value < 0.05 and an absolute log2 fold change > 1 (Type 1D analysis, excluding *h.legs*). Only OGs meeting these criteria in the same direction across all three species were included in the “3-way Conserved Core.” These intersections were visualized using the ggVennDiagram package.

The statistical significance of the observed overlaps was validated using a Hypergeometric test (Representation Factor), accounting for the total size of the orthologous background (8924 OGs). For visualization of the conserved expression profiles, VST-normalized counts for these core OGs were Z-score scaled within each species to generate a consensus heatmap of high-synchrony candidates.

To map the biological processes governing the moult cycle, Over-Representation Analysis (ORA) was performed on the sets of OGs identified in the conserved core for each phase. Enrichment was conducted using the clusterProfiler^62^ and GO.db^63^ packages, utilizing standardized “Preferred Names” from the OrthoDB mapping as the cross-species vocabulary. Significant Gene Ontology (GO) terms for Molecular Function (MF) and Biological Process (BP) were identified for each phase using a Hypergeometric test with Benjamini-Hochberg p-value correction (padj < 0.05).

To characterize regional physiological specializations, we performed within-phase functional enrichment on DEGs identified from two specific anatomical axes (Type 4 analysis): Hind Legs vs. Front Legs (to map the functional polarity of the biphasic transition) and Head vs. Front Legs (to identify conserved cephalic trajectories).

To visualize the successive physiological “waves,” significant terms were filtered and grouped into biologically relevant domains (e.g., chitin metabolism, mineral/ion transport, and proteolysis/recycling). Terms that reached statistical significance in at least two species simultaneously were prioritized for visualization, ensuring that the resulting functional narrative reflected robust, cross-species physiological transitions.

To identify conserved co-expression modules across the isopod lineage, a dual biclustering strategy was applied to the ComBat-seq batch-corrected VST counts of the top 2000 high-variance OGs. This analysis used the ComplexHeatmap package with k-means partitioning (k=8 for rows) to group orthologues with synchronized expression profiles across all species, body parts, and phases.

Two distinct grouping strategies were implemented: 1. Fully Unsupervised Biclustering: Samples and genes were partitioned independently using k-means (4 column clusters, 8 row clusters) to determine if the conserved physiological signal outweighed species-specific or technical variance. 2. Phase-Structured Clustering: Columns were explicitly partitioned by the biological factor “moult phase” to resolve the specific functional modules associated with each stage of the cycle.

The resulting modules were functionally categorized based on their constituent OGs and assigned to the “Universal Moult Toolkit” to quantify the temporal and spatial orchestration of the physiological wave.

Anatomical transcriptomic decoupling was quantified by performing within-phase differential expression contrasts between the front and back legs (Type 4 analysis). For each of the four moult phases, the total number of significant DEGs (|log2FC| > 1, padj < 0.05) between the two body parts was calculated for each species.

To test the “Biphasic Chimera” model, the significance of the transcriptomic surge during the intramoult phase was evaluated using a Wilcoxon rank-sum test, comparing the number of body part-specific DEGs in the intramoult phase against the combined baseline of all other phases. The magnitude of this decoupling was expressed as a fold increase relative to the intermoult baseline. Conserved drivers of anatomical asymmetry were identified by intersecting these DEG lists across the three species to isolate a robust suite of genes (e.g., calcium pumps, cuticle proteins, and immune factors) governing the physiological desynchrony inherent to the biphasic moult. All of the code used in this study is available at https://github.com/IdanSheizaf/TranscriptomicChimera/

## Results and Discussion

### Cyclical Transcriptomic Progression

We initially set out to provide a global characterization of the molecular characteristics of the moulting cycle in terrestrial isopods. We used the transcriptomic profiles of the four selected body parts, among the four study species across the four phases of the moult cycle to characterize the global transcriptomic dynamics of the moult cycle and determine to what extent stage transitions are continuous within individual terrestrial isopod species. We performed independent Principal Component Analysis (PCA) on the VST-normalized count matrices of *A. maculatum* (Figure 1a), *P. laevis* (Figure 1b), and *A. officinalis* (Figure 1c). For each species, the datasets were filtered to focus on the top 2000 high-variance transcripts. This unsupervised analysis revealed a continuous, circular trajectory in transcriptomic space. The phases of the moult cycle form distinct clusters that map onto the two-dimensional representation as a ‘transcriptomic clock.’

**Figure 1:**
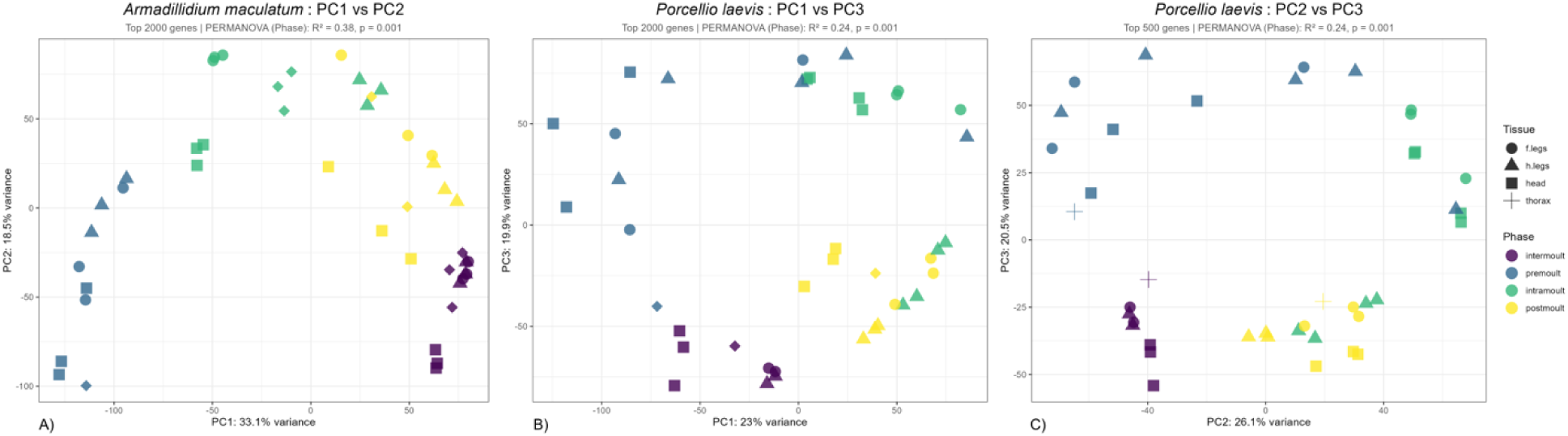
Species-specific transcriptomic clocks of the moult cycle across three terrestrial isopod families. Principal Component Analysis (PCA) of RNA sequencing of four different body parts at four different phases of moulting. The analysis is based on the top 2,000 high-variance transcripts within each species: **(a)** *A. maculatum*, **(b)** *P. laevis*, and **(c)** *A. officinalis*. Samples are colored by physiological moult phase: intermoult (Purple), premoult (Blue), intramoult (green), and postmoult (Yellow). Different body parts are represented by shapes: front legs (circles), hind legs (triangles), head (square), thorax (diamond). For each species, the transcriptomic trajectory exhibits a continuous, circular progression (a ‘transcriptomic clock’), where the physiological moult phase is the primary driver of global variance (PERMANOVA, p < 0.001, with R^2 values ranging from 0.24 to 0.38). During the intramoult, hind leg samples consistently decouple from the remaining anterior body parts (front legs, heads, and thoraces) of the same individuals, clustering with the postmoult samples.

All three species exhibit a rhythmic trajectory where the physiological moult phase represents the primary driver of global transcriptomic variance. This relationship was validated by Permutational Multivariate Analysis of Variance (PERMANOVA, p < 0.001, with R² values ranging from 0.24 to 0.38 across the three species). A key finding from these individual clocks is the spatial separation of body parts during the Intramoult phase. Specifically, hind leg samples from the intramoult phase consistently group with the postmoult samples, whereas the front legs, heads, and thoraces of the same individuals remain clustered with or close to the active ecdysial stages.

These individual trajectories demonstrate that the moult cycle is not characterized by disjointed transcriptomic shifts, but rather by a continuous and highly coordinated physiological loop. The clustering of the hind leg samples with postmoult samples during the intramoult phase suggests that the posterior half of the animal has already initiated the post-moult genetic programme while the anterior half is still preparing for ecdysis. This transcriptomic decoupling demonstrates the molecular underpinnings of the biphasic moulting. The similar pattern across three divergent isopod families suggests that the temporal orchestration of the moult cycle is evolutionarily conserved.

### Conserved Driver of Phase Transition

To determine to what extent there is a core genetic programme shared across the Oniscidea and to isolate conserved signals from phylogenetic divergence and technical noise, we combined the transcriptomic data of all three species into a single orthologous matrix consisting of 8924 shared Orthologous Groups (OGs). To account for species-specific batch effects that could obscure the biological signal, we applied ComBat-seq to the raw count matrix, using species as the batch factor while preserving the moult phase signal. We then performed a global PCA on the top 2000 high-variance OGs (Figure 2a).

**Figure 2:**
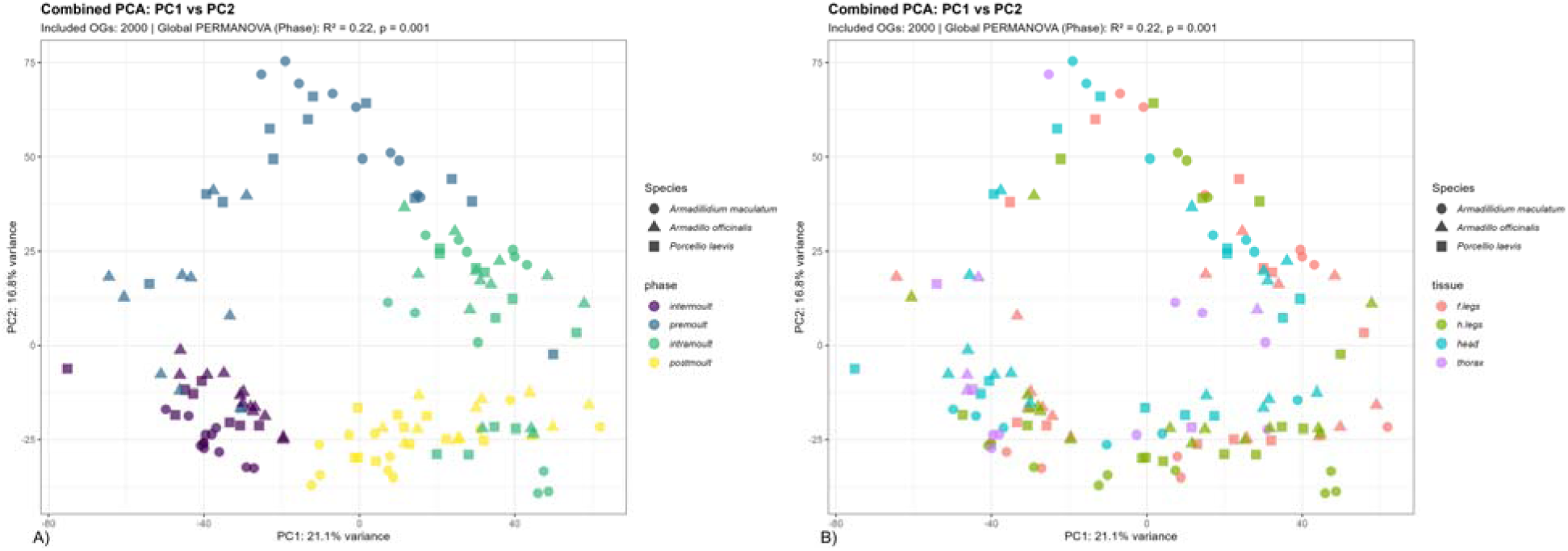
Global integrated transcriptomic alignment and anatomical decoupling. Integrated global PCA of the unified cross-species expression matrix consisting of 2000 shared Orthologous Groups (OGs) across *P. laevis*, *A. maculatum*, and *A. officinalis*, showing a unified developmental trajectory (PERMANOVA, p < 0.001, R^2 = 0.22) **(a)** Samples coloured by physiological moult phase. **(b)** Samples coloured by anatomical body parts.

This analysis revealed that the first two principal components, PC1 and PC2 (which represent the shared transcriptional trajectory), place all three species along a common path (PERMANOVA p < 0.001, R² = 0.22). This alignment confirms that despite more than 100 million years of divergence^37^, terrestrial isopods share a conserved programme, which follows along the different moult phases. Additionally, when colored by body part (Figure 2b), the global analysis shows that all the intramoult hind leg samples across all species cluster within the postmoult groups of other body parts, confirming that the aforementioned decoupling is a conserved feature across the suborder.

To identify the key biological drivers of this shared trajectory, we extracted the OGs with the highest loading scores along PC1 and PC2 (Table S1). PC1 loadings are dominated by genes associated with cuticle synthesis and chitin binding, such as *cuticle protein CP14.6-like*^64^ and *AM1274-like*^65^. In contrast, PC2 loadings capture genes associated with regulatory timing and metabolic homeostasis, including *methyl farnesoate epoxidase-like* (MFE, an enzyme involved in MF metabolism)^66^, *Lysozyme*, participating in chitin breakdown^13,67^, and *peroxidase*^67^, alongside a distinct set of structural genes including cuticle protein CP1499-like^68^, which plays a role in calcification, and collagen alpha-1(XI) chain-like^69^, involved with extra-cellular matrix composition, that load in the opposite direction.

The alignment of all three species on a single trajectory shows that the molecular coordination of the moult cycle has been strictly preserved during the evolution of terrestrial isopods. The partitioning of structural elements across both PCs, with PC2 additionally capturing regulatory/metabolic timers, suggests a multi-layered genetic programme. While PC1 represents the core progressive assembly of the new exoskeleton, PC2 likely represents the temporal coordination and segregation of specific cuticle types alongside hormonal triggers. This modular organization may provide the physiological flexibility needed to coordinate the two-stage biphasic moult on land.The Transcriptomic Chimera and Biphasic Asymmetry

Following up on the observed molecular divergence between the anterior and posterior segments of the body during the biphasic moult, we performed differential gene expression analysis between the front legs and hind legs within the same individuals across the four phases of the moult cycle. This allowed us to measure transcriptomic asymmetry, defined as the number of differentially expressed genes (DEGs) between the two body parts, and test if the biphasic moult is characterized by a molecular desynchronization.

Our results show a significant increase in transcriptomic asymmetry specifically during the Intramoult phase (Figure 3). During the Intermoult phase, transcriptomic symmetry is largely maintained, with an average of only 51 DEGs detected between front and hind legs (across the three species). However, during the Intramoult phase, the number of body part-specific DEGs increases to an average of 1,715, representing a 33.6-fold increase relative to the Intermoult baseline This increase is statistically significant (Wilcoxon Rank Sum Test, p = 0.009). The transcriptomic decoupling in space indicates that the organism is deploying two different gene expression programmes for body parts of the same type, with the anterior and posterior halves executing distinct physiological programmes, in what we call a ‘transcriptomic chimera’. Analysis of the OGs that drive this asymmetry (Table S2) indicates that the decoupling is organized into two functional modules. The front legs, which remain in a pre-ecdysial state, exhibit a significant enrichment in genes involved in cuticle degradation and recycling. This includes a synchronized set of *V-type proton ATPase* subunits (subunits B, C, F, H, and S1) that are required for acidifying the extracellular space to facilitate cuticle digestion^70^, alongside proteases such as Trypsin, and larval cuticle proteins. Additionally, the front legs show strong upregulation of UDP-glucuronosyltransferase 2C1-like, which may locally regulate active ecdysteroid levels to delay anterior ecdysis^71^.

**Figure 3:**
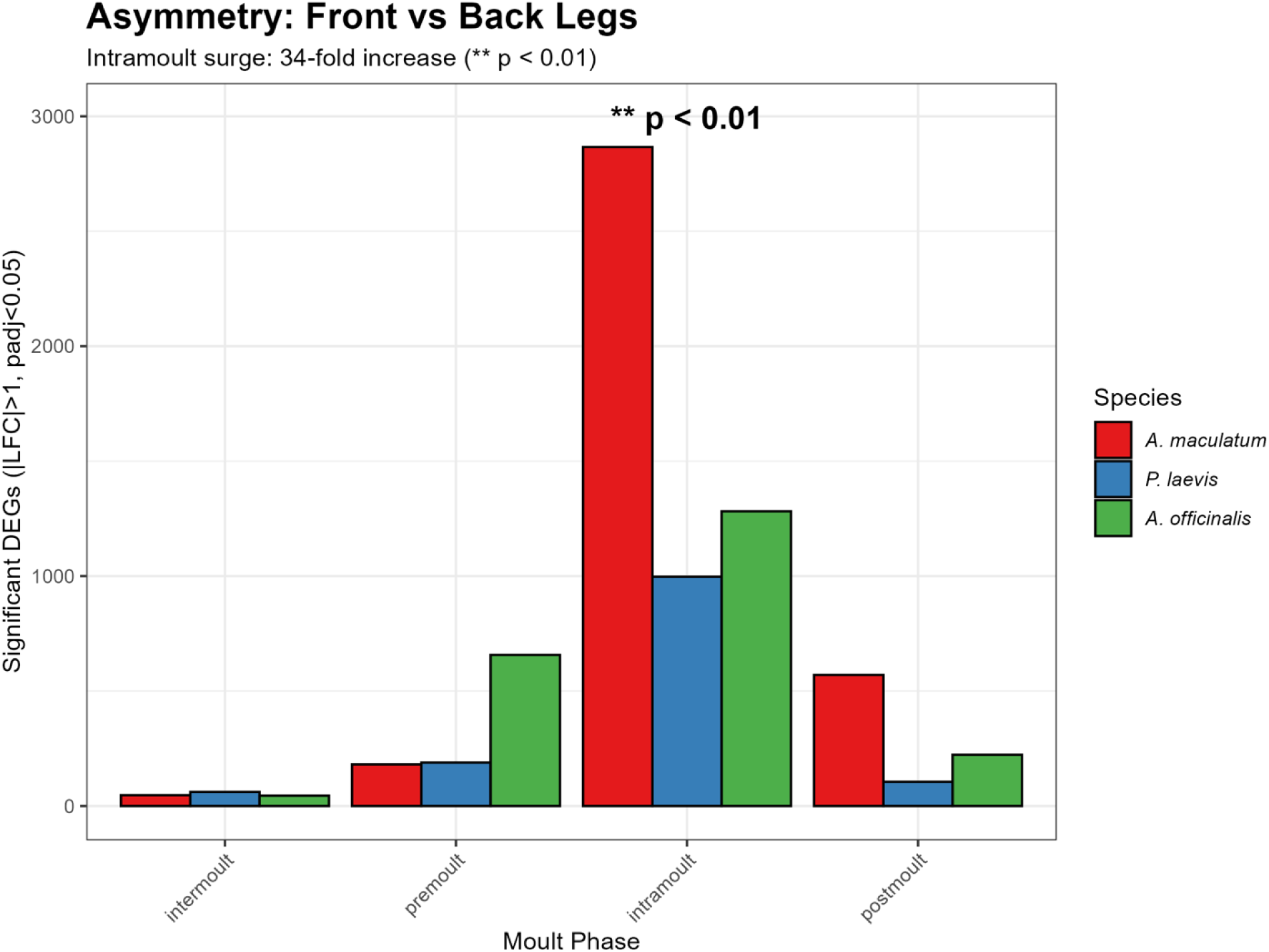
Rhythmic surge in transcriptomic asymmetry during the intramoult phase. Anatomical transcriptomic decoupling quantified via pairwise differential gene expression contrasts between front and hind limbs within individual animals across the four moult cycle phases. The bar plot details the distribution of body part-specific differentially expressed genes across the three species. Under physiological baseline conditions (intermoult), transcriptomic symmetry is maintained, yielding an average of only 51 DEGs. During the Intramoult interval, asymmetry surges to a mean of 1,715 DEGs, a statistically significant 33.6-fold increase (Wilcoxon Rank Sum Test, $p = 0.009$).

In contrast, the hind legs, which have completed exuviation, show a strong bias for genes involved in immune defense and extracellular matrix stabilization). This includes Lysozyme, Anti-lipopolysaccharide factors (ALFs), and protein spaetzle-like (which collectively form a protective barrier against pathogens^72,73^), which may be driven upstream by the upregulation of the Nuclear factor NF-kappa-B subunit, of the innate immune response^74,75^ . Cellular and matrix reconstruction in the posterior segments is further characterized by the upregulation of collagen alpha-1(XI) chain-like, unconventional myosin-X-like, associated with vesicular transport and cell remodelling, and ECM-anchoring proteins like Spondin and hemicentin-1-like. This structural reorganization is coordinated by the regulatory nuclear hormone receptor FTZ-F1 beta^76,77^ and fuelled by solute carrier family 2 glucose transporters^78^.

This spatial decoupling demonstrates that the behaviourally evident biphasic moult is driven by a coordinated molecular division of labour. By staggering ecdysis, terrestrial isopods avoid the vulnerability associated with simultaneous whole-body shedding. The active anterior half allows the animal to maintain mobility to seek humid microhabitats and avoid predators, while the posterior half initiates cuticle mineralization. Furthermore, because a newly shed cuticle is soft and highly susceptible to bacterial infection and physical damage, the posterior segment expresses a protective suite of immune genes (*Lysozyme*, *ALFs*). This localized immune response suggests that the animal prioritizes defense in its most vulnerable region, which is a key physiological adaptation for survival in microbe-rich soil and leaf-litter environments. Note that while we have compared only terrestrial isopod species, the biphasic moulting is characteristic of all isopods, including marine and freshwater species. We do not have enough data to speculate on the original source of this pattern in the isopod common ancestor, but it must continue to provide an advantage in terrestrial environments, as detailed above, evidenced by its conservation.

### Conserved Expression Modules and Toolkits

We next aimed to identify co-expressed gene groups that exhibit similar expression profiles across all three species. We therefore performed a dual-clustering analysis on the shared OGs. We generated a comprehensive set of heatmaps using the top 2000 high-variance OGs (Figures S1-S2, Table S5). Figure 4 presents only the top 100 high-variance OGs to allow visual and biological clarity. We conducted both a fully unsupervised biclustering analysis (Figure 4a, S2) and a phase-split clustering analysis (Figure S1, S3), using the ComplexHeatmap package in R. The unsupervised biclustering grouped samples primarily by their moult phase rather than by species or body part identity, confirming that the conserved transcriptomic signal of the moult is strong enough to overcome phylogenetic differences.

**Figure 4:**
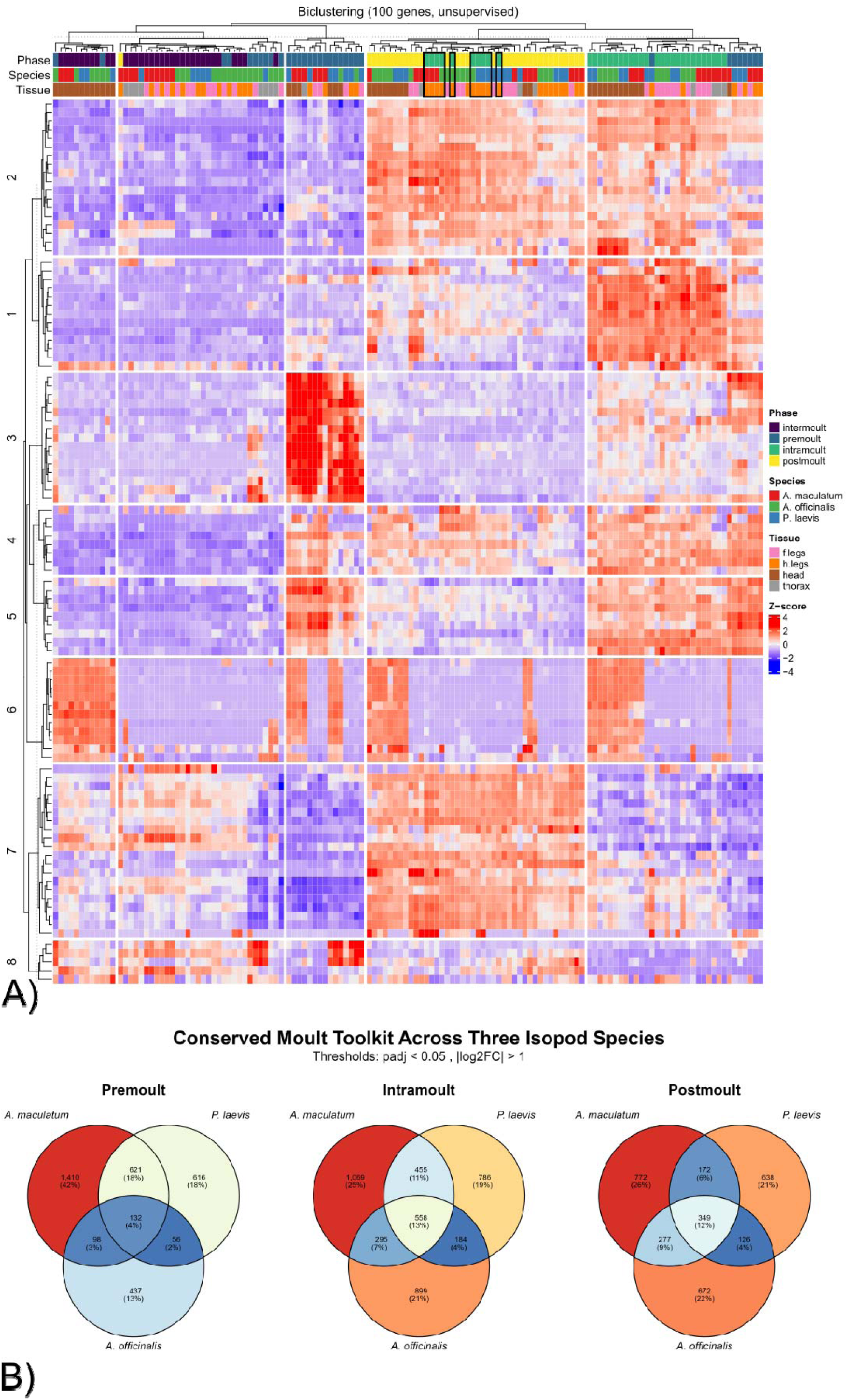
Conserved co-expression modules and core molecular toolkit of the isopod moult. Consensus transcriptomic orchestration resolved via biclustering and directional overlaps. (a) Heatmap representing Z-score scaled, VST-normalized expression profiles of the top 100 high-variance OGs. The unsupervised biclustering partitions samples primarily by physiological stage rather than species or body part and separates genes into 8 synchronized modules (Modules 1–8; annotated in Table S3), mapping structural cascades (Module 1, chitin/cuticle), endocrine triggers (Module 5, methyl farnesoate epoxidase), and post-ecdysial immune resetting (Module 7, lysozyme/ALFs). (b) 3-way Venn diagram showing the overlap of significantly regulated OGs relative to the intermoult baseline in all three species (excluding hind legs to prevent early-transition bias). The intersection identifies the conserved core ‘molecular toolkit’ driving the transitions. The statistical significance of the overlap was validated via a Hypergeometric test against the 8924 OG background.

By organizing the shared transcriptome into eight major conserved expression modules (Table S3), we were able to map the functional progression of the cycle. Module 1, representing the intramoult climax and anatomical decoupling, is dominated by structural components such as *chitin-binding domain* proteins and *cuticle proteins*. This module is highly active in all intramoult body parts except for the hind legs samples (marked by black frames), which fits the observation that the posterior half transitions early. Module 2 represents the transition from intramoult to postmoult and is enriched for *flexible cuticle proteins* and *nucleolin*, which may coordinate the transition from active shedding to early matrix stabilization. Module 3, which peaks in premoult, is biased toward the head and includes signalling and extracellular matrix genes like *mastermind-like domain-containing protein 1*^79^ and *adhesive plaque matrix protein-like*. Modules 4 and 5 capture pre-ecdysial priming in premoult, showing enrichment for neuromuscular proteins such as *myosin* and ecdysis signalling elements like *MFE*. Module 6 represents cephalic sensory and neuroendocrine regulation, showing enrichment for *crustacean hyperglycemic hormones (CHH)* and *opsins*, which may coordinate the systemic timing of the cycle. Module 7 represents late-phase homeostasis and is dominated by *Lysozyme* and *ALFs*, which may protect the newly shed animal from pathogens. Finally, Module 8 represents a more heterogeneous metabolic group that lacks clear phase-locking, possibly reflecting general housekeeping roles.

To define the core ancestral genes of the moult, we intersected the significantly regulated DEGs (each phase vs. the Intermoult reference) across all three species (Figure 4c; Table S4). To avoid the confounding effect of early-transitioning body parts, the hind leg samples were excluded from this analysis. This intersection identified a core “molecular toolkit” of conserved orthologues. For example, during the Intramoult phase, the toolkit includes structural drivers such as *Chitin binding domain* (OG: 47160at6657, 28581at6657) and *Cuticle proteins* (OG: 30821at6657), as well as immune factors like *Anti-lipopolysaccharide factor* (OG: 67125at6657), *Lysozyme* (OG: 42138at6657), and *C-type lectins* (OG: 10139at6657). Metabolic and regulatory genes, including *V-type proton ATPase* subunits (OG: 36044at6657, 51402at6657) and *Ecdysteroid kinase-like* (OG: 69370at6657), were also conserved.

Grouping the transcriptome into these co-expression modules reveals that the complex process of ecdysis is driven by a series of synchronized waves of gene expression. The phase-based clustering in the unsupervised analysis indicates that the physiological state of the moult is a primary determinant of gene expression, overriding species boundaries. However, some caveats must be noted: because functional annotations are largely based on sequence homology with other crustaceans (often based originally on holometabolous insects), the precise role of many of these orthologues in terrestrial isopods remains speculative. For instance, the presence of neuromuscular genes in Module 4 may indicate active structural remodelling of leg muscles to facilitate ecdysis, but direct functional validation (e.g., via RNA interference) is required to confirm this. Similarly, the specialized neuroendocrine signature of Module 6 highlights the head’s role in system-wide coordination, but the exact triggers for phase transitions remain to be fully characterized.

### Anatomical Decoupling and Central Hormone Regulation

To determine if transcriptomic shifts are linked with changes in anatomical regions over time, we mapped functional enrichment waves across body parts. We specifically contrasted locomotory body parts (hind legs vs. front legs) to characterize the biphasic transition and compared cephalic tissues (head vs. front legs) to identify conserved regional specializations.

#### Physiological Polarity of the Biphasic Moult: Hind Leg – Front Leg Asymmetry

We contrasted the transcriptomic profiles of hind legs and front legs across the four phases of the moult cycle. During the Intramoult phase, we observed a distinct functional polarity between the two segments (Tables S6-S7). In the hind legs, we identified enrichment in *monoatomic ion transport* (GO:0006811; Figure 5a) and *calcium ion binding* (GO:0005509; Figure 5b), driven by genes encoding calcium pumps, sodium-potassium ATPases, and carbonic anhydrases. In parallel, the hind legs showed enrichment in *proteolysis* (GO:0006508) and *serine-type endopeptidase activity* (GO:0004252), whereas the front legs remained enriched in cuticle-degrading enzymes.

**Figure 5:**
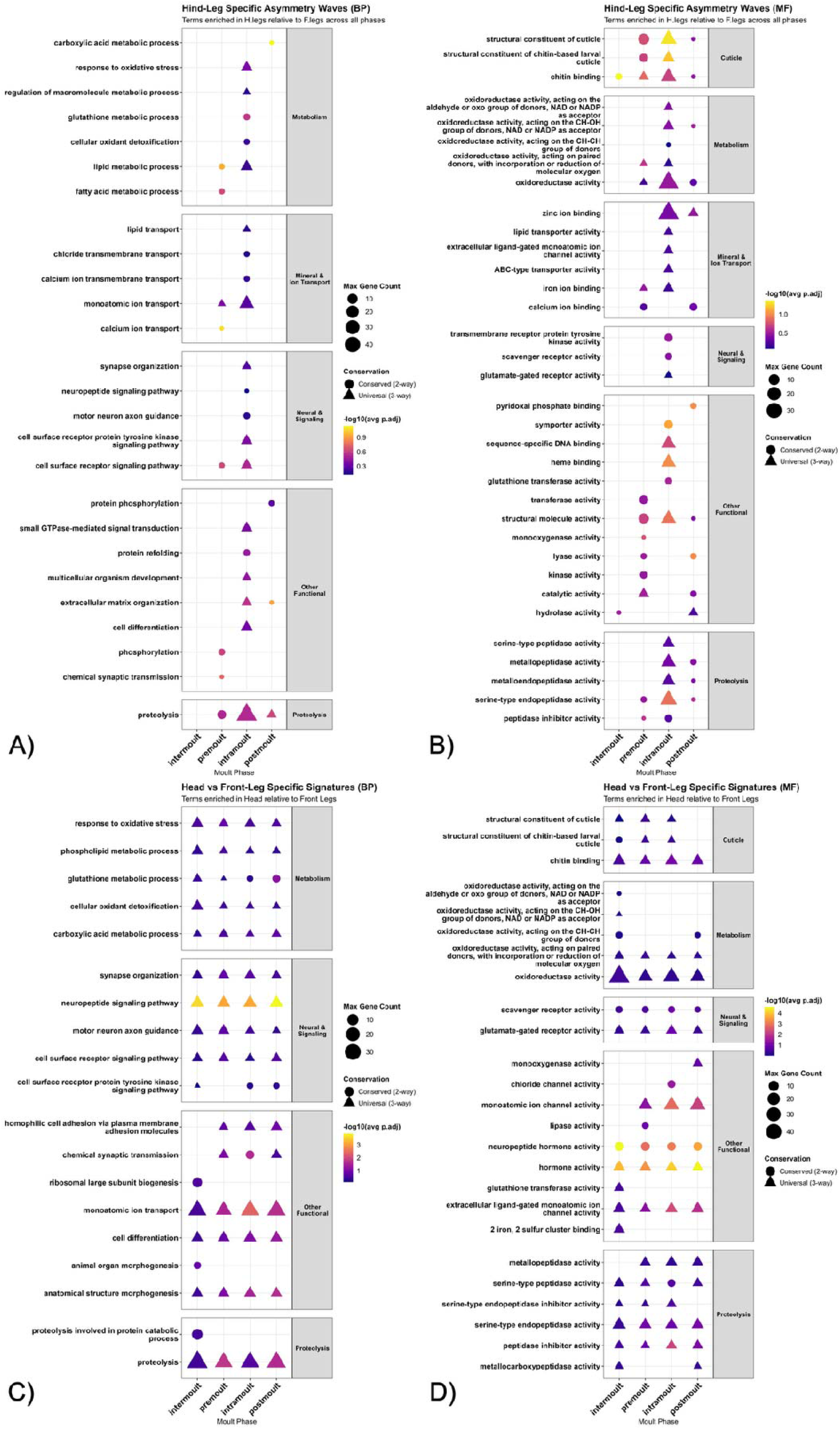
Body part-specific functional enrichment waves and regional specializations. Functional Gene Ontology (GO) enrichment profiling mapping body part-specific trajectories (Hypergeometric test, Benjamini-Hochberg corrected padj < 0.05; terms showing a similar pattern – up- or down-regulation – in at least two species are prioritized). **Head vs. front Legs (a-b):** Represents the stable neuroendocrine regulatory system of the head. Cephalic tissues exhibit a highly stable enrichment of neuropeptide signalling pathway (GO:0007218) and chemical synaptic transmission (GO:0007268) (a), alongside hormone activity (GO:0005179) and neuropeptide hormone activity (GO:0005184) (b) across all moult phases, contrasting with the high structural/transcriptomic turnover observed in the limbs. **Axial asymmetry - hind legs vs. front legs (c-d):** Captures the functional polarity of the biphasic switch. During the Intramoult interval, hind legs (posterior) display strong enrichment of monoatomic ion transport (GO:0006811) (c) and calcium ion binding (GO:0005509) (d), driven by calcium pumps, ATPases, and carbonic anhydrases coordinating rapid post-ecdysial mineralization. Conversely, the front legs (anterior) remain enriched in cuticular digestion, chitin breakdown, and steroid deactivation, facilitating the pre-ecdysial programme.

These functional analyses support the physiological decoupling inherent to the biphasic moult. The enrichment of ion transport and calcium binding in the hind legs suggests that this segment is actively mobilizing and depositing calcium to calcify the new cuticle, while the front legs are still digesting the old exoskeleton. In terrestrial environments, where calcium cannot be absorbed directly from surrounding water, isopods must rely on temporary internal storage (such as sternal calcium deposits). The localized activation of calcium-binding and transport systems in the posterior half immediately after ecdysis represents a critical adaptation for rapid calcification on land, minimizing the time the animal remains soft and vulnerable to desiccation and predation.

### Conserved Cephalic Stability: The head neuro-endocrine trajectory

In contrast to the functional shifts observed in the locomotory body parts, the head transcriptome exhibits a stable functional profile throughout the moult cycle (Tables S8-S9). When comparing the head to the front legs, we identified a suite of biological processes that remain enriched in the head across all stages. This profile is dominated by *neuropeptide signaling* (GO:0007218) and *chemical synaptic transmission* (GO:0007268) (Figure 5c), alongside a consistent enrichment in *hormone activity* (GO:0005179) and *neuropeptide hormone activity* (GO:0005184) (Figure 5d).

These findings suggest that the cephalic tissue does not undergo the wide transcriptomic turnover observed in the limbs. Instead, it operates as a stable neuroendocrine regulatory system. By maintaining a steady expression of neuropeptides and hormones (such as *crustacean hyperglycemic hormone* and *ecdysis-triggering hormone receptor-like* genes), the cephalic region coordinates the systemic timing of ecdysis across the rest of the body. This centralized control is likely essential for ensuring that the posterior and anterior ecdysial events are properly spaced, preventing premature shedding of the anterior cuticle before the posterior half is sufficiently calcified to support the animal.

### The Conserved Candidate Atlas for Isopod Ecdysis

Finally, we wanted to identify the core molecular drivers of the terrestrial isopod moult. We therefore analysed the intersection of the conserved Venn core (genes significant in at least 2 species, p < 0.05) with the major driver genes of the shared PCA trajectory (PC1 and PC2). Filtering identified a broad suite of candidate orthologues (Table S10). Figure 6 presents a selection of 20 high-impact genes from the candidate gene atlas. These candidates were selected by ranking genes based on their PC loading importance and prioritizing those with identified biological roles in literature, while omitting uncharacterized proteins (which are retained in Table S10 for future functional validation). These 20 genes can be organized into three primary biochemical modules:

**Figure 6:**
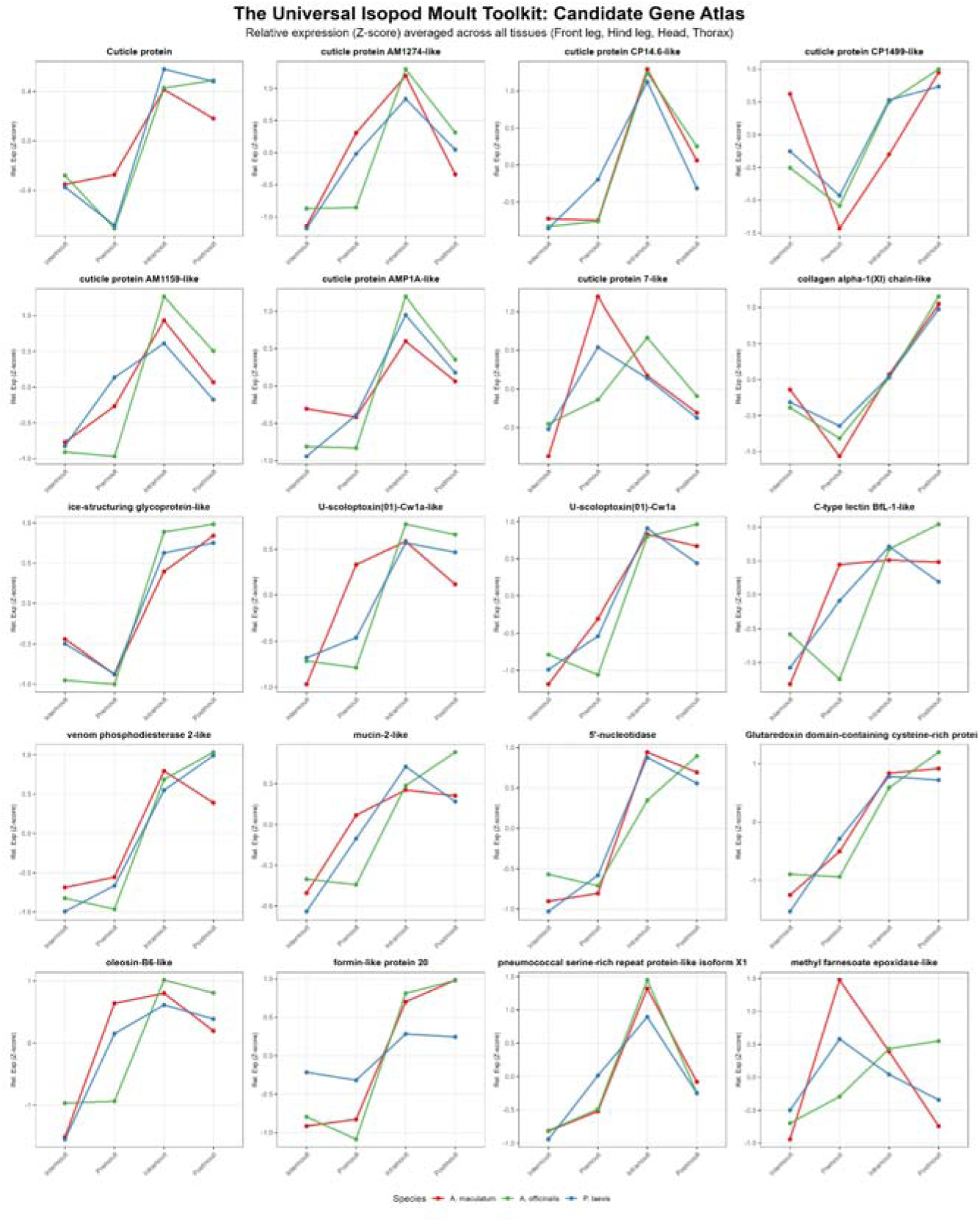
Conserved candidate gene atlas for terrestrial isopod ecdysis. Gene expression grid displaying Z-score scaled expression profiles of 20 candidate orthologues across species and moult phases. Candidates were selected by intersecting the 3-way Venn conserved core with the top PCA driver loadings (PC1/PC2), ranked by importance, and filtered to exclude uncharacterized proteins. The atlas highlights three major functional modules: Structural Biosynthesis and Matrix Remodelling: Co-expressed cuticle proteins (AM1274-like, CP14.6-like, CP1499-like, AM1159-like, AMP1A-like, cuticle protein 7-like), collagen alpha-1(XI) chain-like, and ice-structuring glycoprotein-like peaking during active synthesis (Intramoult/Premoult). Protective and Defensive Adaptations: Defensive factors peaking during vulnerable phases, featuring U-scoloptoxin(01)-Cw1a-like and U-scoloptoxin(01)-Cw1a (Intramoult), and C-type lectin BfL-1-like (Premoult). Terrestrial Homeostasis and Endocrine Regulators: Adaptations to land, including mucin-2-like (lubrication during exuviation to prevent desiccation-induced trapping), 5’-nucleotidase, glutaredoxin domain-containing cysteine-rich protein, oleosin-B6-like, formin-like protein 20, pneumococcal serine-rich repeat protein-like, and methyl farnesoate epoxidase-like (MFE).

### Structural Biosynthesis and Matrix Remodelling

This module is dominated by genes loading on PC1, reflecting the structural requirements of exoskeleton synthesis. It features a co-expressed group of *cuticle proteins* (including *AM1274-like*, *CP14.6-like*, *CP1499-like*, *AM1159-like*, *AMP1A-like*, and *cuticle protein 7-like*), which exhibit peak expression during the intramoult and premoult phases. This matrix assembly is supported by *collagen alpha-1(XI) chain-like* and *ice-structuring glycoprotein-like* (reclassified here as a cuticle-remodelling factor). While not shown in the 20-gene grid, the broader toolkit (Table S10) includes *Chitin synthase (chs-2)*, *Chitin-binding domain* proteins, and *ADAMTS* metalloproteinases that coordinate procuticle polymerization.

### Protective and Defensive Adaptations

This module contains defensive genes that peak during the most vulnerable phases of ecdysis. *U-scoloptoxin(01)-Cw1a-like* and *U-scoloptoxin(01)-Cw1a* peak sharply during the Intramoult phase. This is complemented by *C-type lectin BfL-1-like*, which peaks during premoult, and *venom phosphodiesterase 2-like*, which peaks during postmoult. The broader toolkit (Table S10) also contains *Lysozyme*, which peaks during the postmoult homeostatic reset.

### Terrestrial Homeostasis and Endocrine Regulators

This module highlights physiological adaptations to land and hormonal timers. *Mucin-2-like* peaks during the Intramoult phase, providing lubrication. Other genes peaking during this transition include *5’-nucleotidase*, *Glutaredoxin domain-containing cysteine-rich protein*, *oleosin-B6-like*, *formin-like protein 20*, and *pneumococcal serine-rich repeat protein-like*. The regulatory programme is further marked by *methyl farnesoate epoxidase-like (MFE)*, an enzyme that regulates juvenile hormone analogue levels.

The expression profiles of these 20 candidates demonstrate how structural synthesis, defence, and hormonal regulation are temporally coordinated. While most candidates show synchronized expression patterns across all three species, some transcriptomic divergence is observed in *A. officinalis*, where certain profiles do not align with those of *A. maculatum* and *P. laevis*. This divergence is consistent with the greater phylogenetic distance of the *Armadillo* lineage compared to the more closely related *Armadillidium* and *Porcellio* genera.

From an evolutionary perspective, several of these candidates are directly linked to the challenges of terrestrialization. Because ecdysis in terrestrial environments lacks the physical support of water, the high expression of *Mucin-2-like* during the Intramoult phase likely provides critical lubrication to prevent the soft cuticle from adhering to the old shell during exuviation, which can lead to fatal trapping. Furthermore, the sharp peak of *U-scoloptoxin(01)-Cw1a-like* during Intramoult suggests a specialized role in protecting the soft, uncalcified posterior exoskeleton from soil-dwelling pathogens and opportunists before the protective cuticle has fully hardened. The synchronization of *MFE* and *Alpha carbonic anhydrase* further highlights the coordination of endocrine timers with calcium mobilization, demonstrating how the ancestral crustacean moult programme was modified to support survival on land.

## Conclusions

In this study, we integrated comparative transcriptomics across three phylogenetically divergent terrestrial isopod species to map the genetic orchestration of the biphasic moult cycle. Our analysis demonstrates that individual species trajectories follow a continuous, clock-like progression through transcriptomic space, driven primarily by the physiological phase of the moult. By constructing a unified orthologous matrix, we identified a conserved transcriptional trajectory that aligns these species, showing that the core timing and structural modules of the moult are highly conserved. Crucially, we characterized the molecular signature of the biphasic switch, revealing a surge in transcriptomic asymmetry during the intramoult interval. This spatial decoupling, where the posterior half has transitioned into post-ecdysial hardening while the anterior remains in pre-ecdysial digestion, defines a molecular chimera state. Within this state, the front of the animal expresses cuticle recycling machinery while the back activates immune defence and calcium-binding transport systems, illustrating how this morphological asymmetry is directly mirrored in molecular asymmetry. Finally, we identified a core molecular toolkit, highlighting genes such as *Mucin-2-like* and *U-scoloptoxin(01)-Cw1a-like* that represent potential physiological adaptations to the challenges of moulting on land. While functional confirmation of these candidates (e.g., via gene knockdown) is needed, our findings establish a robust genomics framework for understanding how the ancestral crustacean moult programme was employed to support life in terrestrial habitats. Ultimately, this work demonstrates that terrestrial isopods serve as an excellent model to study the dynamics of moulting, offering a powerful system to explore complex spatial and temporal physiological coordination.

## Author Contribution

I.S. experiments, analysis, writing. R.M.W. analysis, discussion, editing. M.R.R. analysis, discussion, editing. A.D.C supervision, discussion, writing.

## Acknowledgments

The authors would like to acknowledge Ariel Bar Lev Viterbo for his critical comments along the way, and the entire Sinergia moulting team for discussions and support.

