## Supplementary figures and images for "The Molecular Programme of the Biphasic Isopod Moult: A Transcriptomic Chimera"

### Figure S1

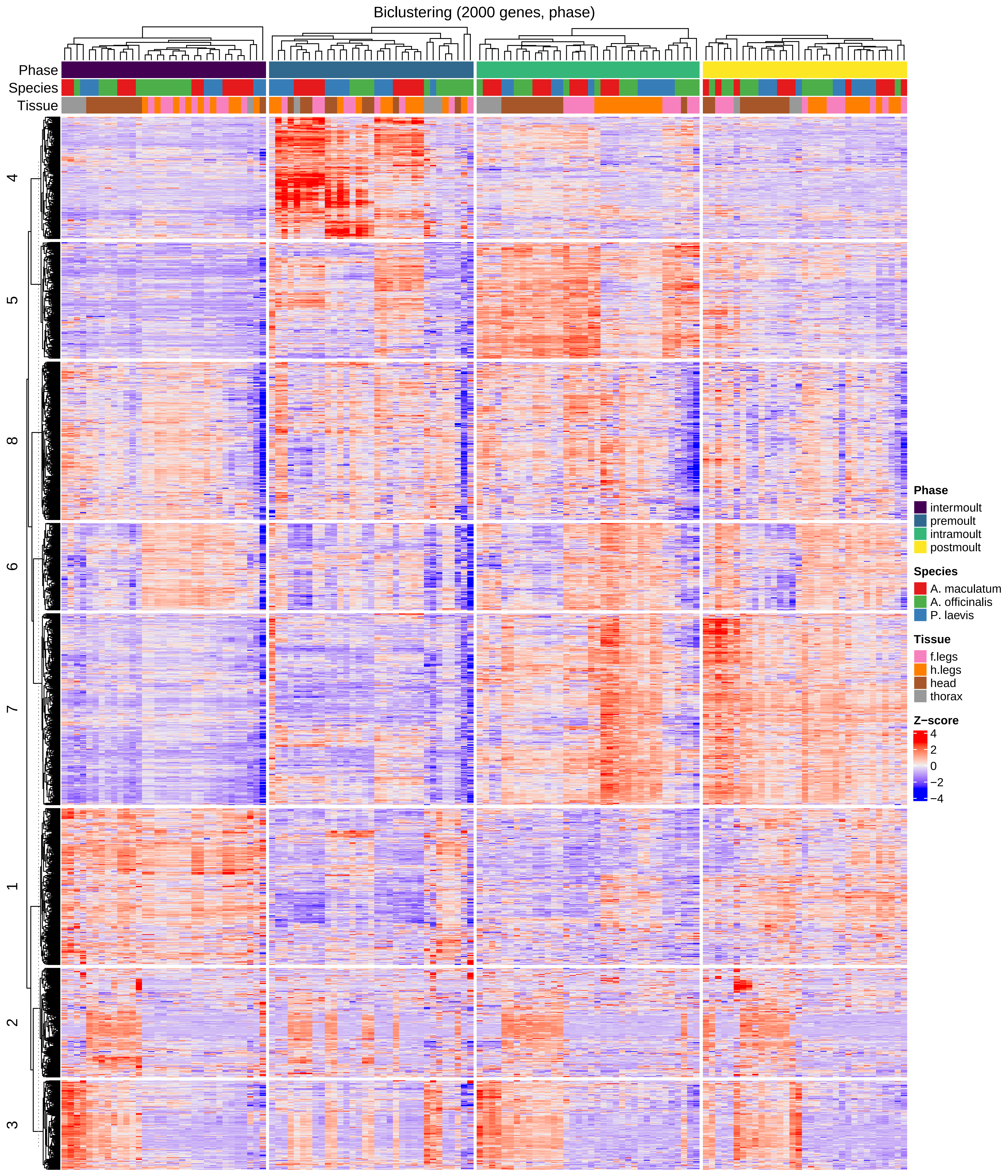

### Figure S2

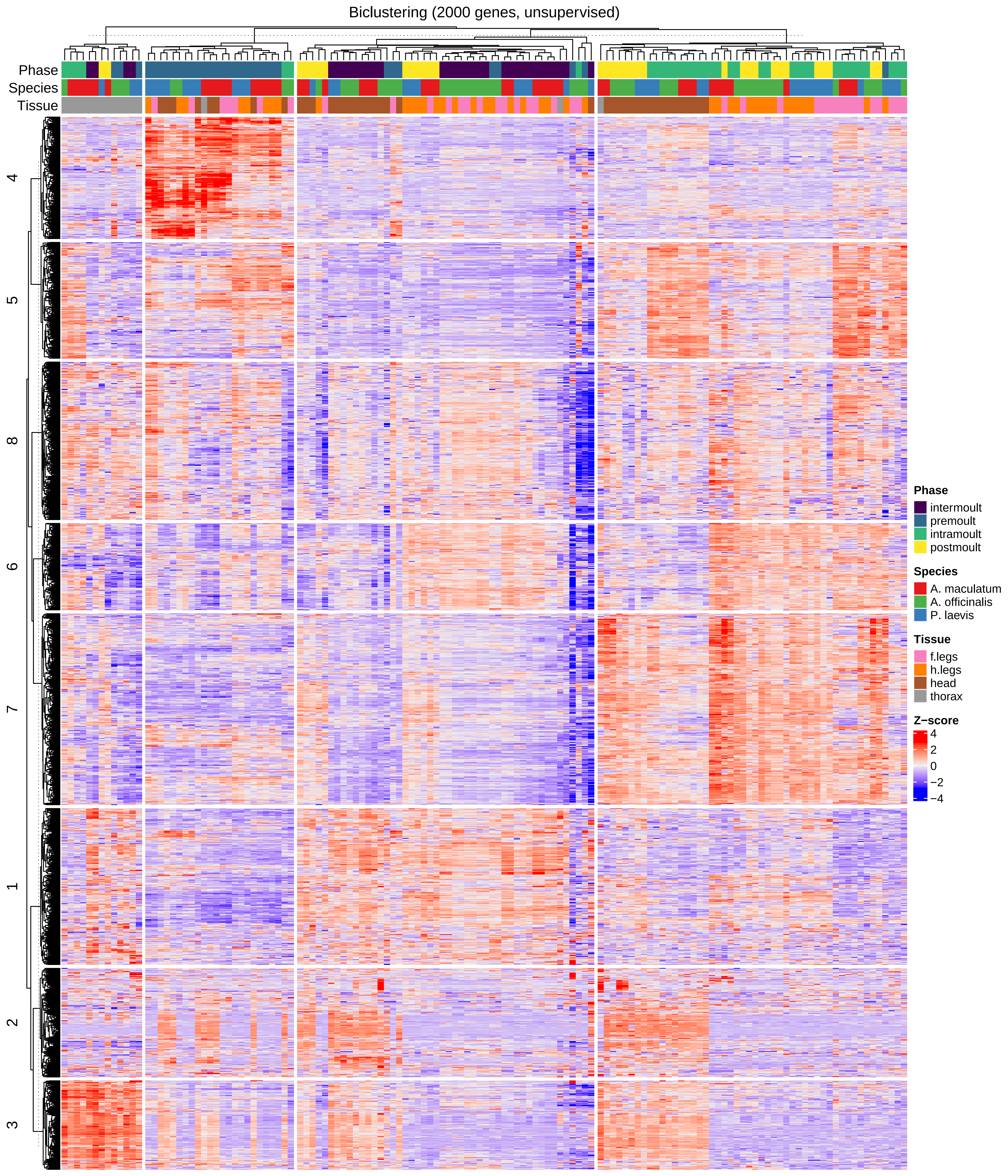

### Figure S3

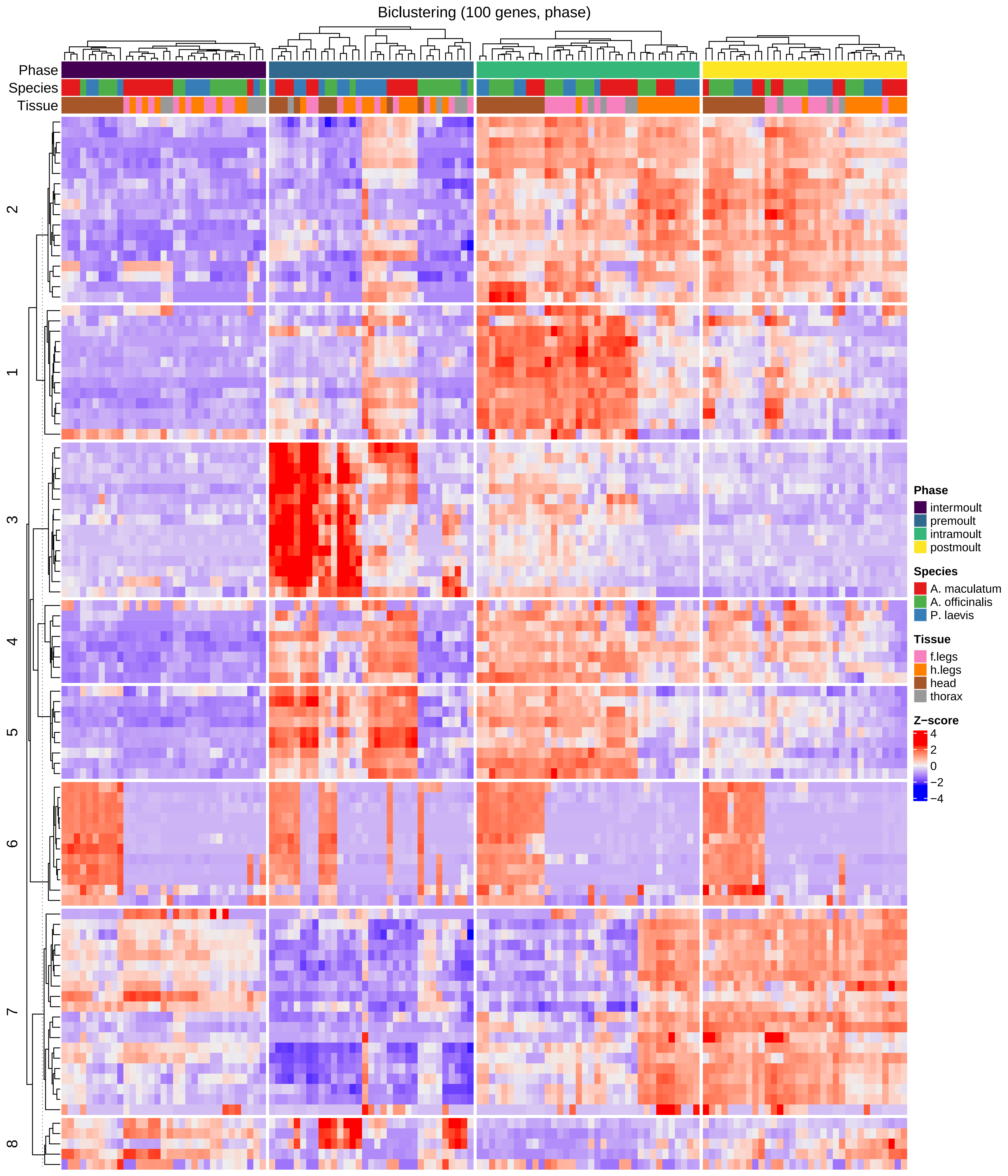
