## Supplementary legends for "The Molecular Programme of the Biphasic Isopod Moult: A Transcriptomic Chimera"

Supplementary Figures and Tables

Figure S1: Phase-structured co-expression heatmap of the top 2000 high-variance OGs, where columns (samples) are explicitly partitioned by biological moult phase. This structured approach resolves the broad transcriptomic waves lock-stepped to the developmental transitions.

Figure S2: Fully unsupervised co-expression heatmap of the top 2000 high-variance OGs. It demonstrates that the global transcriptional state transitions serve as the dominant axis of variance, grouping samples by moult phase across species and tissues.

Figure S3: Phase-structured co-expression heatmap of the top 100 high-variance Ogs, where columns (samples) are explicitly partitioned by biological moult phase. This structured approach resolves the broad transcriptomic waves lock-stepped to the developmental transitions.

Table S1: Global PCA driver genes and loadings.

Spreadsheet listing the top 100 Orthologous Groups (OGs) driving the integrated global PCA trajectory under ComBat-seq batch correction. Details include ODB_OG identifiers, preferred gene names, individual loadings along PC1 and PC2, and calculated Importance Scores (absolute loading multiplied by explained variance). Features prominent cuticle-binding proteins (CP14.6-like, AM1274-like), calcification drivers (CP1499-like), matrix stabilizers (collagen alpha-1(XI)), and enzymatic timers (methyl farnesoate epoxidase, lysozyme, peroxidase).

Table S2: Conserved drivers of anatomical asymmetry.

Complete list of conserved orthologs driving front-to-hind leg asymmetry during the Intramoult interval. It highlights the division of labor between the pre-ecdysial front legs (upregulated in acidifying V-type ATPase subunits B, C, F, H, S1, Trypsin, and UDP-glucuronosyltransferase) and post-ecdysial hind legs (upregulated in antimicrobial lysozyme, ALFs, spaetzle, matrix hemicentin-1, Spondin, and glucose transporters).

Table S3: Co-expression modules of the top 100 high-variance OGs.

Spreadsheet categorizing the 100 high-variance OGs into 8 discrete co-expression modules corresponding to Figure 4a-b. Includes functional classifications (e.g., chitin metabolism, neuromuscular, neuroendocrine regulation, post-moult homeostatic resetting).

Table S4: Conserved molecular toolkit of the isopod moult.

Spreadsheet listing the universal 'molecular toolkit' orthologs identified in the 3-way Venn diagram intersection (Figure 4c) for Premoult, Intramoult, and Postmoult transitions relative to the Intermoult baseline. Includes detailed log2 fold changes and adjusted p-values for each species.

Table S5: Co-expression modules of the top 2000 high-variance OGs.

Spreadsheet containing module membership of the top 2,000 OGs. It defines 8 major co-expression clusters: (1) Intermoult cluster, (2) Head-specific cluster, (3) Thorax/Head cluster, (4) Premoult-specific cluster, (5) Intramoult-specific cluster, (6 & 8) Exoskeletal assembly cluster, and (7) Late-phase/Postmoult reset genes.

Table S6: Biological Process consensus enrichment for locomotory tissue asymmetry.

Consensus Biological Process (BP) enrichment terms for the Front vs. Hind Legs comparison across all four phases. Details min/avg/max adjusted p-values and count of species in which the term reached significance.

Table S7: Molecular Function consensus enrichment for locomotory tissue asymmetry.

Consensus Molecular Function (MF) enrichment terms for the Front vs. Hind Legs comparison across all four phases.

Table S8: Biological Process consensus enrichment for Head vs. Front Legs.

Consensus Biological Process (BP) enrichment terms for Head vs. Front Legs comparison, mapping stable neuroendocrine processes.

Table S9: Molecular Function consensus enrichment for Head vs. Front Legs.

Consensus Molecular Function (MF) enrichment terms for Head vs. Front Legs comparison, mapping hormone and receptor activity.

Table S10: Extensive list of conserved candidate orthologs.

Comprehensive registry of conserved candidate genes extracted by intersecting orthologs of the conserved Venn diagram core with PC1 and PC2 PCA drivers. Details include ancestral ODB_OG IDs, preferred gene names, PC loadings, importance scores, primary expression phase, and biochemical classifications (e.g., Structural/Cuticle, Immunity/Protection, Lubrication/Desiccation).
